# The Genotype and Phenotypes in Families (GPF) platform manages the large and complex data at SFARI

**DOI:** 10.1101/2024.02.08.579330

**Authors:** Liubomir Chorbadjiev, Murat Cokol, Zohar Weinstein, Kevin Shi, Chris Fleisch, Nikolay Dimitrov, Svetlin Mladenov, Simon Xu, Jake Hall, Steven Ford, Yoon-ha Lee, Boris Yamrom, Steven Marks, Adriana Munoz, Alex Lash, Natalia Volfovsky, Ivan Iossifov

## Abstract

The exploration of genotypic variants impacting phenotypes is a cornerstone in genetics research. The emergence of vast collections containing deeply genotyped and phenotyped families has made it possible to pursue the search for variants associated with complex diseases. However, managing these large-scale datasets requires specialized computational tools tailored to organize and analyze the extensive data. GPF (Genotypes and Phenotypes in Families) is an open-source platform (https://github.com/iossifovlab/gpf) that manages genotypes and phenotypes derived from collections of families. The GPF interface allows interactive exploration of genetic variants, enrichment analysis for *de novo* mutations, and phenotype/genotype association tools. In addition, GPF allows researchers to share their data securely with the broader scientific community. GPF is used to disseminate two large-scale family collection datasets (SSC, SPARK) for the study of autism funded by the SFARI foundation. However, GPF is versatile and can manage genotypic data from other small or large family collections. Our GPF-SFARI GPF instance (https://gpf.sfari.org/) provides protected access to comprehensive genotypic and phenotypic data for the SSC and SPARK. In addition, GPF-SFARI provides public access to an extensive collection of *de novo* mutations identified in individuals with autism and related disorders and to gene-level statistics of the protected datasets characterizing the genes’ roles in autism. Here, we highlight the primary features of GPF within the context of GPF-SFARI.

## Introduction

The substantial reduction in sequencing costs has made it feasible to generate whole-exome or whole-genome sequences for large family collections. This breakthrough allows for the direct observation and analysis of genetic variants across the entire frequency spectrum, spanning from common to rare and *de novo* mutations. What’s more, large-scale sequencing in genetic studies provides researchers with the ability to pinpoint the causal variant itself rather than relying on genetic markers that merely point to a genomic region harboring these variants. Over the past five years, numerous sequencing studies involving hundreds to thousands of families have significantly advanced our understanding of the genetic underpinnings of various phenotypes. Notably, this approach has yielded remarkable insights, particularly in the context of childhood disorders that have a substantial impact on an individual’s reproductive ability, such as autism spectrum disorder (ASD) and congenital heart disorders (CHD), where *de novo* mutations were confirmed to play a pivotal causal role. However, the complexity and scale of data generated by sequencing large family collections surpasses that of traditional genotyping studies. As a result, there is a pressing need to devise specialized methodologies and tools for the efficient management, analysis, and dissemination of this vast and intricate dataset.

The development of such tools is an actively evolving field with numerous examples. The GnomAD Browser [1] provides access to variants from extensive whole-genome and whole-exome datasets, encompassing approximately 750,000 exomes and 80,000 whole genomes. It includes a diverse human population sample and provides a powerful interface for access to the population/summary variants but excludes individual genotypes. Similarly, the Bravo variant server (https://bravo.sph.umich.edu) allows access to population variants identified from nearly one million individuals in the NIH’s TOPMed program [2], spanning various heart, lung, and blood disorders and controls. DeNovoDB [3] is a repository for *de novo* variants identified in individuals, particularly children with autism and related neurodevelopmental disorders. While it offers a user-friendly interface for querying and downloading these variants, it lacks information about transmitted/Mendelian variants in these individuals. Notably, these tools neither handle family data directly nor integrate transmitted variants within families with identified *de novo* variants. Moreover, they lack interactive analysis tools and are specifically tailored to specific datasets, posing challenges for adaptation elsewhere.

GPF (Genotypes and Phenotypes in Families) is a platform that can handle millions of individuals’ genetic variants and deep phenotypic data. We developed GPF to manage the data related to the Simons Simplex Collection (SSC), a collection of ∼2,800 families, each of which has one child affected with autism [4]. The platform proved extensible and flexible enough to accommodate the larger SPARK (Simons Foundation Powering Autism Research) collection, a growing collection of ∼100,000 individuals with autism and their families [5, 6], as well as other collections supported by SFARI. Our GPF instance, GPF-SFARI, enables researchers to interactively conduct complex queries for variant selection and perform genotype-phenotype association and gene-set variant enrichment analyses using the SSC and SPARK datasets. Due to the sensitive nature of these datasets, users need permission to access the full extent of SSC and SPARK. Still, GPF-SFARI has two publicly accessible components for unregistered users: the “Sequencing *de novo*” dataset contains published *de novo* mutations for six developmental disorders in typically developing children; the “Autism Gene Profile” contains summary statistics for all human genes, including the number of variants of various types in the SSC, SPARK, and “Sequencing *de novo*” datasets, and gene properties relevant to autism.

While GPF is developed specifically for the SFARI collections, it is designed so that other researchers may create their own GPF instances and analyze their genotypic and phenotypic data using the provided tools and visualizations. Here, we will describe the general features of GPF and the datasets within the GPF-SFARI instance. Then, we will describe the query, enrichment, and phenotype tools GPF provides in the context of GPF-SFARI. Finally, we discuss how users can use GPF to analyze and share their genotypic and phenotypic data.

## Results

### General features of GPF

The Genotypes and Phenotypes in Families (GPF) system manages genotypes and phenotypic measures of individuals from collections of families. Genotypes of various types identified through different technologies (whole-exome, whole-genome, etc.) and genotyping tools are imported into the system. Separately, phenotypic measures are imported. The system then provides an intuitive interface for exploring, jointly analyzing, and securely sharing the genotypic and phenotypic data.

GPF is designed to accommodate a diverse range of family structures (Figure 1a). The basic family type is the nuclear family, with two parents and their children. Nuclear families (especially the trios with two parents and one child) are the main family types used in the study of *de novo* mutations. GPF also supports complex multi-generational families like the ones frequently used in linkage analysis of Mendelian phenotypes. Additionally, to accommodate the needs of case-control studies, the system supports families comprised of single individuals. GPF uses pedigree files to capture the family relationships and basic phenotypic information, such as gender, affected status, and the indication of the individual proband.

**Figure 1.**
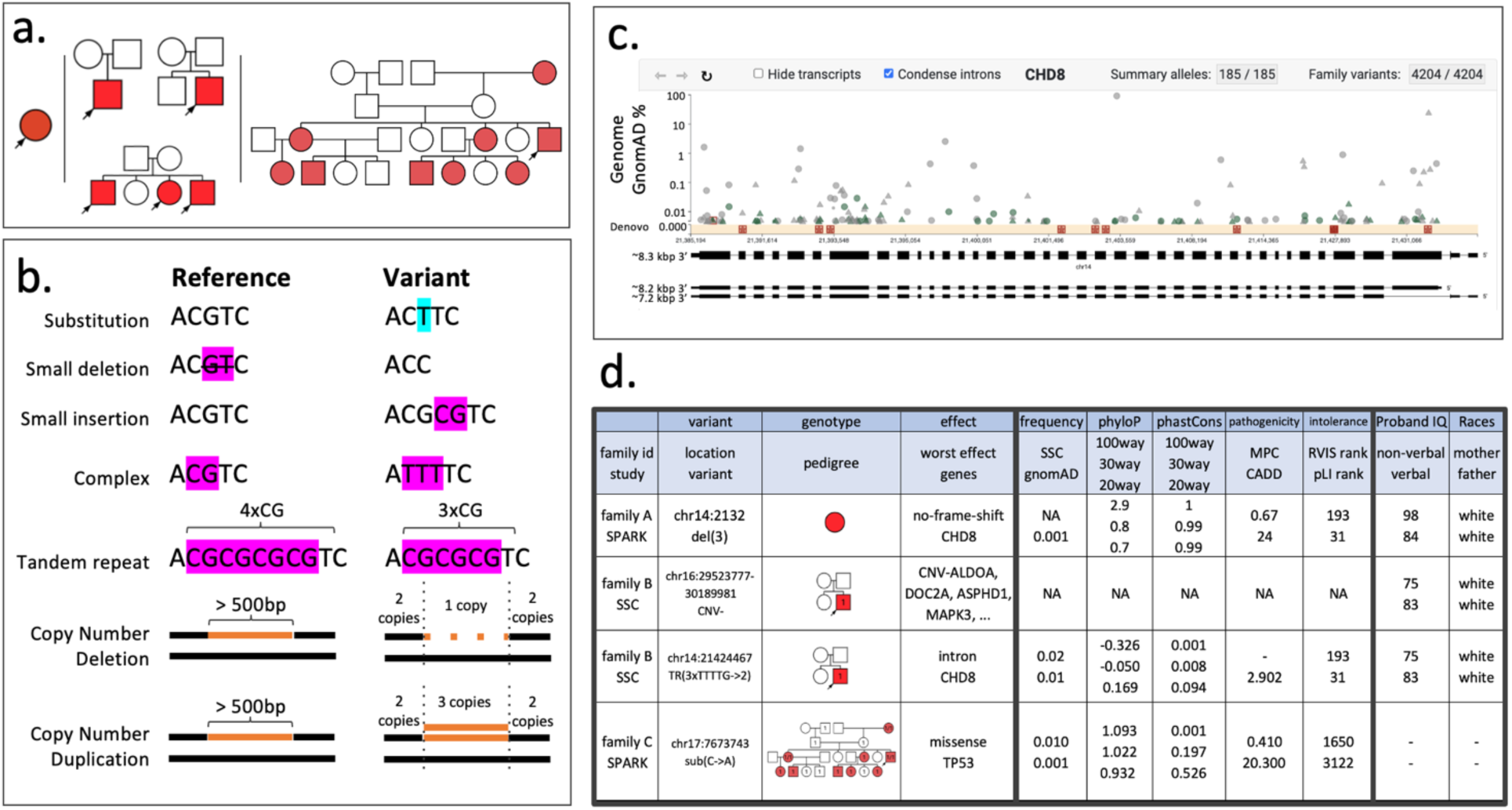
General Features of GPF. (a) GPF supports various family structures represented as pedigree diagrams, where parent-child relationships are shown with lines, males and females with squares and circles, respectively, and phenotypes with colors. In this panel, red indicates individuals diagnosed with autism, and white indicates individuals not diagnosed with autism. Shown are a family with a single individual (left), three nuclear families (two parents and their children) (middle), and a complex multi-generational family (right). The arrows indicate the probands in the families. (b) GPF supports various genetic variant types represented relative to a reference genome, including substitutions, short insertions and deletions, copy number variants, and tandem repeats (or microsatellites). (c) The gene view component shows the population (or summary) variants identified in a gene of interest. The example shows the variants in the *CHD8* gene. The gene view represents the genomic location of the variants on the X axis in the context of the gene isoforms and the variant frequencies on the Y axis. Stars, triangles, and circles are likely-gene disruption (LGD), missense, and synonymous variants, respectively. Red squares indicate *de novo* variants. (d) The family variants view shows four *CHD8* variants segregating in four families. The family variants are represented as rows in a table organized in three sections of columns. The first section shows the family (the family id/study column), the variant (the location/variant column), the segregation pattern (the pedigree column), and the variant’s predicted effect on the protein-coding genes (the worst effect/genes column). In the segregation patterns, numbers indicate the alternative alleles segregating in the family, e.g., an individual with ‘1’ is heterozygous for the first alternative allele, and an individual with ‘1/1’ is homozygous. The second section shows the GPF’s extensive genomic annotations assigned to each variant. These annotations may include frequencies of the variant in the reference populations (e.g., gnomAD [1]), conservation measures (e.g., phyloP [10]and phastCons [11]), pathogenicity scores (e.g., MPC [12] and CADD [13]), and gene intolerance scores (e.g., RVIS [14] and pLI [15]). The third section shows relevant phenotypic measures of the proband or their family members, demonstrating the ability of GPF to integrate phenotypic information and the genetic variants segregating in the family.

GPF supports a wide range of variant types, each representing distinct genetic alterations (Figure 1b). The simplest variant is a substitution, where a single nucleotide at a position differs from the nucleotide at the same position in the reference genome. Small deletions and insertions (indels) involve the removal or addition of 1 or a few nucleotides in comparison to the reference genome. Complex variants manifest when a short sequence is replaced by another sequence, potentially of a different length. Tandem repeats are short sequences of nucleotides repeated multiple times, prone to changes in the number of repeats during cell division, and very polymorphic in human populations. Copy number deletions and duplications encompass instances where a large (i.e., exceeding 500 base pairs) segment of DNA is either removed or duplicated. These variant types can be identified through various technologies. For example, single nucleotide variants (SNVs), indels, and copy-numbers variants (CNVs) can be detected using methods like Whole Exome Sequencing (WES) and Whole Genome Sequencing (WGS), while single nucleotide polymorphisms (SNPs), and CNVs can also be identified through array hybridization.

GPF further categorizes alleles based on their inheritance patterns within a family. Alleles exhibiting a Mendelian transmission pattern are designated as ‘Mendelian.’ An allele present in a child but not inherited from either parent is categorized as ‘*de novo*.’ Lastly, when an allele should logically be passed on to a child according to Mendelian genetics but is not, it falls under the classification of ‘omission.’

In GPF, population variants for a specific gene are visually represented through the Gene View, a widely adopted method for visualizing genetic variants. In the Gene View, variants are depicted as symbols on a scatter plot, providing information about their genomic locations (x-axis), frequencies (y-axis), and types (symbol shapes). Figure 1c shows the gene view for 185 alleles in *CHD8*, a chromodomain helicase DNA-binding protein associated with autism [7, 8]. GPF’s Gene View is interactive, allowing users to select subsets of the gene variants by type, location, or frequency and obtain the families where the selected variants segregate in the Family Variants View.

The Family Variants View offers users insight into genetic variants segregation in families. Figure 1d exemplifies this by presenting information on four *CHD8* family variants organized into three sections of columns. The first section outlines essential details about the variant, including its familial context, segregation pattern within the family, and a prediction of its impact on protein-coding genes. The second section shows in-depth genomic annotations assigned to each variant during the import process. These annotations include a range of data, such as variant frequencies in reference populations like gnomAD [9], conservation metrics like phyloP [10] and phastCons [11], pathogenicity scores like MPC [12] and CADD [13], and gene intolerance scores like RVIS [14] and pLI [15]. The third section contains phenotypic measures, highlighting GPF’s capacity to integrate information about family members’ phenotypes and the genetic variant’s segregation within the family structure.

GPF organizes its data into genotype and phenotype studies. The genotype studies consist of a set of genotypes for the individuals from a set of families. Similarly, phenotype studies consist of phenotypes from individuals from a set of families. Phenotypes in GPF are further structured into instruments or diagnostic instruments applied to the study’s individuals. Each instrument includes a set of specific measures. A genotype study can be linked with a phenotype study, forming a cohesive connection between genetic and phenotypic information. Datasets, in turn, are composite groupings of studies and previously defined datasets. The studies grouped into datasets can contain similar data types for different sets of families or different data types for the same families. GPF allows for a detailed description of each of its studies and datasets. This flexible framework enables users to work on individual studies or to analyze multiple studies or one or across all datasets jointly.

### GPF at SFARI

The Simons Foundation Autism Research Initiative (SFARI) aims to improve the understanding, diagnosis, and treatment of autism spectrum disorders (ASD) by supporting and funding innovative scientific research. Simon Simplex Collection (SSC) [4] and Simons Foundation Powering Autism Research (SPARK) [5, 6], the largest collection of families with autism, are the two main initiatives of SFARI.

SSC encompasses roughly 2,800 simplex families, each including a single child with autism, the affected child’s parents, and, for most families, one or more unaffected siblings. SSC has comprehensive individual phenotypic profiles, especially for the affected probands. The phenotypes are based on the application of diagnostic instruments, including autism diagnostic tools like ADI-R, ADOS, measures of core features of autism and intellectual ability, and family history [4]. SFARI has funded several large projects for generation array hybridization genotypes and whole-exome and whole-genome sequencing data from all SSC individuals. These data have been used by many research groups to transform the study of autism genetics, demonstrating the large contribution of *de novo* mutations to autism and establishing a process for identifying autism genes using *de novo* variants that has produced hundreds of high-confidence genes [16–20].

Following the success of SSC, SFARI initiated SPARK, a growing collection of self-registered families with autism, which currently has ∼90,000 families or approximately 300,000 individuals [5, 6]. SPARK includes both multiplex and simplex families, accommodating a wide range of family structures. The consent forms allow researchers to re-contact participants of interest, and through the recontacting process, there is a growing list of phenotypic measures of interest. SFARI has funded the generation of whole-exomes from ∼120,000 SPARK individuals at Regeneron and whole-genomes from an additional ∼30,000 individuals at the New York Genome Center (NYGC).

Through their data-sharing policy, SFARI requires researchers to return/deposit the data they have generated through the sequencing datasets back to SFARI. These returned data include genotypes of various types, CNV, SNV, indels, microsatellites, etc., both *de novo* and inherited. Part of SFARI’s mission is to distribute these data back to the scientific community. However, these data are large and complex, and most are sensitive and require protection through strict authorization processes.

SFARI uses GPF to share the data, allowing outside (non-SFARI) researchers to explore and analyze the large and complex datasets of genotypes in deeply phenotyped families. The user can also use the genetic and phenotypic data in GPF to select materials distributed by SFARI, such as DNA, cells, or tissues from the individuals in the SSC and SPARK collections. We’ll refer to the GPF deployed at SFARI as GPF-SFARI (https://gpf.sfari.org/).

A subset of the data in the GPF-SFARI is freely accessible to everyone. The rest comprises sensitive genotypic and phenotypic data that require a rigorous authorization process to grant access to qualified researchers. SFARI has implemented such a process through their SFARI Base infrastructure that governs the access to the sensitive data within GPF and, generally, all the data and materials managed by SFARI. Figure 2a shows the major data components of the GPF-SFARI. The “Sequencing *de novo*” component is a dataset that contains freely accessible lists of *de novo* variants found in the SSC and SPARK collections, in other collections with autism, as well as *de novo* variants identified in children with five additional developmental disorders. The two largest datasets contain the protected genotypic and phenotypic data related to the SSC and SPARK collections. The “Gene Profile” component integrates freely accessible summary information of the data from the three components organized by gene. The Gene Profile also contains external information relevant to studying the genes’ roles in autism. Below, we provide more details for each of the four major components of GPF-SFARI.

**Figure 2.**
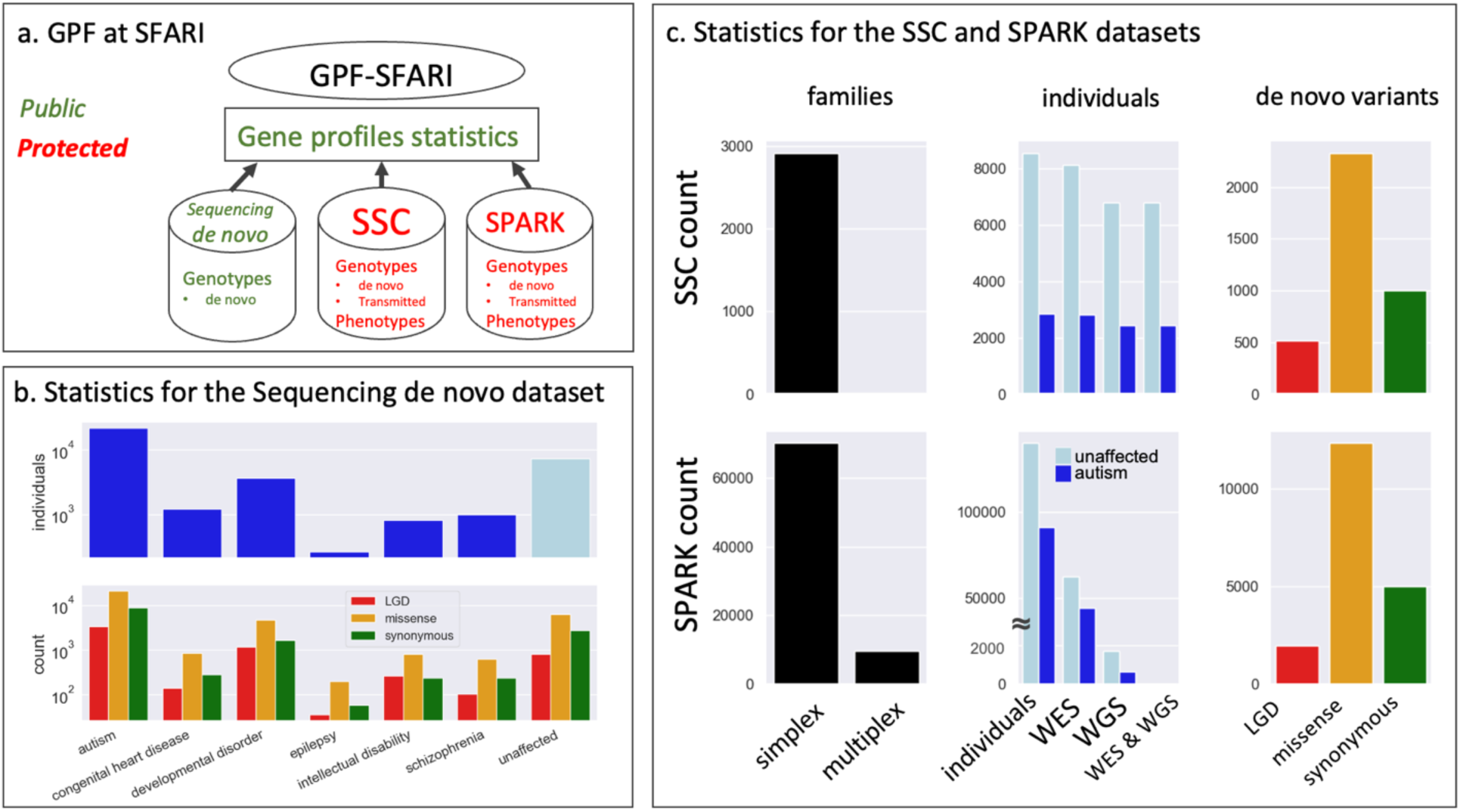
GPF-SFARI. (a) We organized the data in the GPF instance deployed at SFARI (GPF-SFARI) into three large datasets. Two datasets, SSC [4] and SPARK [5, 6], are protected under a rigorous authorization process and comprise the *de novo* and transmitted genotypes and phenotypes from the two large collections of families with autism built by SFARI. The third dataset is a collection of publicly available *de novo* variants identified in individuals diagnosed with one of six developmental disorders (autism, schizophrenia, intellectual disability, congenital heart disease, and epilepsy) and in typically developing children labeled as unaffected. In addition, the GPF-SFARI includes a Gene Profile component, which is also publicly accessible. The Gene Profile component represents information relevant to autism for each human gene, including the number of various types of variants in the three datasets. The data in the Gene Profile is organized as one all-genes table (see Table 1) and individual gene pages. The Gene Profile allows unregistered users to see aggregate information from otherwise protected SSC and SPARK datasets. (b) The panel shows the number of individuals (top) and the number of *de novo* coding variants (bottom) included in the Sequencing de novo dataset separately for each diagnosis. The bottom section shows the counts for three types of *de novo* coding variants: *de novo* likely-gene disruption (LGD) variants (blue), *de novo* missense variants (orange), and *de novo* synonymous variants (green). (c) The panel shows the numbers of families (left), individuals (middle), and *de novo* coding variants (right) in the SSC (top) and SPARK (bottom) datasets. The numbers of families are organized by family type: simplex or multiplex. Simplex families are families with only one individual diagnosed with autism, and multiplex families have more than one. The numbers of individuals are split based on the diagnosis: dark blue for individuals diagnosed with autism (affected) and light blue for the ones not diagnosed with autism (unaffected). We show the numbers of individuals with phenotypic data (individuals) and those with genotypes derived from whole-exome (WES), whole-genome (WGS), or both (WES & WGS). Finally, the numbers of *de novo* coding variants are presented separately by the variant effects: LGD, missense, or synonymous.

**Table 1.**
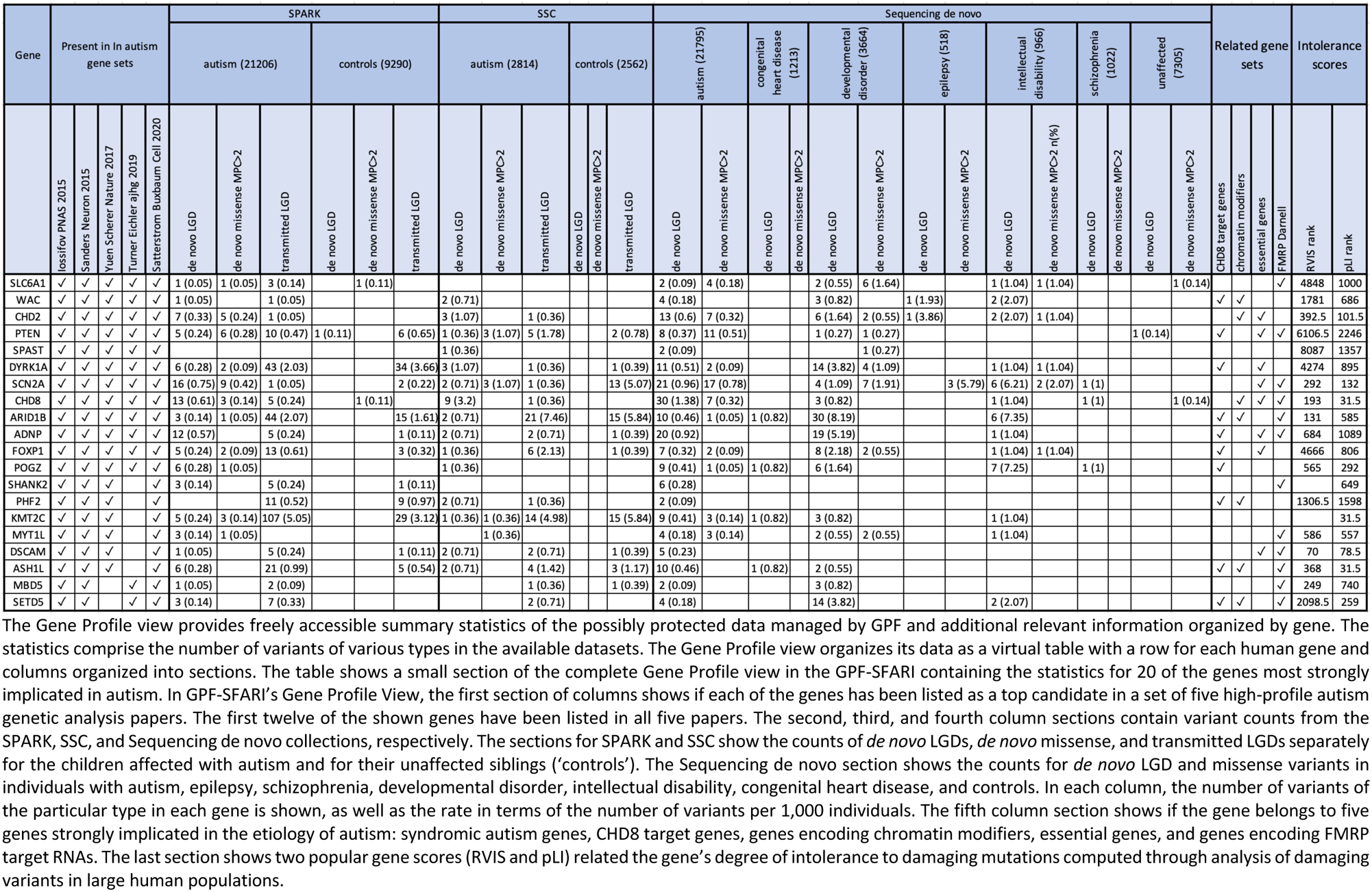
Gene Profile View.

The “Sequencing *de novo*” is a comprehensive resource that compiles lists of *de novo* substitution and short indels associated with six distinct abnormal development disorders and *de novo* variants found in typically developing children. The dataset predominantly comprises data from published research articles, providing a valuable reference for genetic research. Additionally, it incorporates de novo variants obtained from the SPARK collection, some of which are yet to be formally published, expanding the breadth of genetic insights available for analysis. Figure 2b shows the number of children and *de novo* LGD, missense, and synonymous variants included in “Sequencing *de novo*” for each disorder. Some of the studies included in “Sequencing *de novo*” are based on whole-genome sequencing data and contain non-coding de novo variants. The “Sequencing *de novo*” description in GPF-SFARI lists the publications used to build the data set.

SSC and its large sequence data have been around for a while and have been extensively studied. Many groups, including us, have applied different tools to generate genotypes of various types. These genotypes, aggregated and imported in GPF-SFARI, include *de novo* and transmitted substitution and indels from the whole-exome [21–26] and whole-genome [19, 27–29] datasets, *de novo* CNVs called from hybridization array [17, 30] and from the whole-genome data [27], and microsatellites called from the whole-genomes [31]. The SSC dataset in GPF-SFARI also contains the complete phenotypic data associated with SSC, comprising ∼100 phenotypic instruments or ∼10,000 individual measures.

SPARK is a more recent and growing collection, so its processing and analysis are ongoing and incomplete. To date, ∼90,000 families comprising ∼300,000 have self-registered in SPARK, but sequence data (whole-exome and whole-genome) has been generated from only a subset (see Figure 2c). Genotypes have been generated and imported in GPF-SFARI for an even smaller subset [5, 6]. The major group that produces genotypes from the whole-exome sequence data is the SPARK Consortium, which includes the SFARI’s bioinformatics team. The SPARK consortium generates updates of the SPARK genotypes regularly, and currently, GPF-SFARI includes transmitted and *de novo* substitutions and indels from ∼106,000 individuals. The numbers of *de novo* LGD, missense, and synonymous variants are listed in Figure 2c and follow the expected rates per individual. In addition, the SPARK dataset contains ∼400,000 and ∼3,000,000 transmitted LGD and missense variants. NYGC performs genotyping from the whole-genome data in batches, and GPF-SFARI includes transmitted substitutions and indels from ∼10,000 individuals from the first large batch of whole-genomes.

While data within SPARK and SSC is safeguarded, we offer unrestricted access to specific aggregate measures about genes through the ‘Gene Profiles View.’ This view provides a comprehensive list of variant counts with various effects for all genes, along with additional relevant gene properties. For instance, within the ‘Gene Profiles View’ on SFARI, you can find counts of *de novo* likely gene-disrupting (LGD), missense, and synonymous variants, as well as the number of transmitted variants involving *de novo* LGD variants from both SSC and the continuously expanding SPARK collection. Furthermore, the tool shows if genes have been identified as ‘autism’ genes in multiple publications [16–20] and if they belong to relevant gene sets, such as FMRP genes [32], CHD8 targets [7], or chromatin modifiers [21, 25]. Additionally, SFARI’s ‘Gene Profiles View’ presents various measures of gene intolerance, including RVIS [14] and pLI [15] scores.

### GPF power tools

#### Genotype browser

GPF provides users with power tools for interacting with the data. One of these key tools is the Genotype browser, which enables users to create and execute intricate queries for genetic variants. Users can search for variants based on the properties of the variants, the properties of the genes the variants affect, and the phenotypic properties of the individuals carrying the variants (Supplementary Figure 1). The variant properties that can be used in queries include the variant location, type, genes, predicted effects on the protein-coding genes they affect, and genomic scores like phyloP [10], CADD [13], and MPC [12], assigned to variants during the import in GPF.

The properties of genes that can be included in a query are of two types: gene set membership and gene scores. Gene sets are a handy abstraction in many biological contexts, including genes in a pathway, the targets of a transcription factor or RBP, and genes containing a given protein domain. Relevant collections of gene sets can be imported into GPF. For example, in GPF-SFARI, we have imported the GO ontology [33], MSigDB [34, 35], sets of published lists of autism genes, and gene sets implicated in autism etiology (i.e., chromatin modifiers, FMRP targets, embryonically expressed genes). Each of the imported gene sets has a name, and users can select gene sets by name to formulate a query like ‘*search for variants affecting genes involved in apoptosis (a pathway from MSigDB) or genes implicated in autism*.’ GPF provides two alternatives to selecting a predefined gene set: the users can provide their gene set as a list of genes or use the GPF genetic variants to define a gene set as the gene affected by *de novo* LGDs in GPF’s dataset. Gene scores are numbers assigned to genes that describe properties like gene length and degree of intolerance to damaging mutation. The relevant gene scores, obtained from external sources and imported into GPF, can be used to formulate queries like, ‘*find genetic variants that affect genes with RVIS intolerance score larger than 4*’.

Finally, users can search for variants based on the phenotypic properties of the individuals carrying the variants. The phenotypic properties include diagnosis, gender, and the measures of instruments in the phenotypic studies. For example, a user can search for variants in male children with autism with IQs less than 75.

Figure 3a presents the result of a complex query through the Genotype browser: five of the 175 de novo LGD variants in affected males within an FMRP target gene are shown. Users may download the selected variants in a comma-separated file format for further analysis beyond the GPF platform. These downloaded files include the genotypes of all family members, along with a comprehensive set of annotation attributes for all variants.

**Figure 3.**
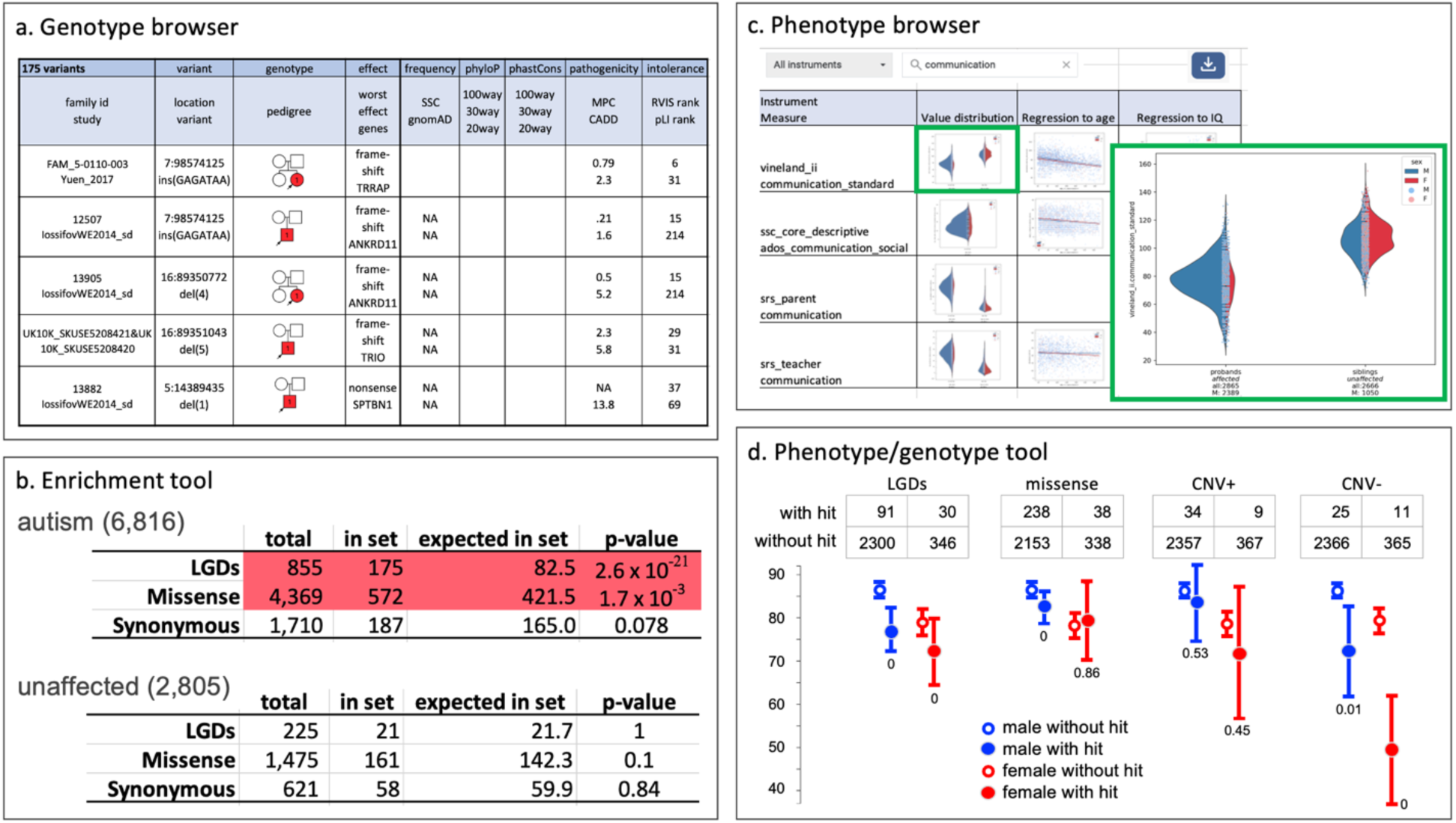
Query and statistical tools at GPF. (a) The Genotype browser allows the user to form complex queries involving the genetic variants’ properties, the affected genes’ properties, and the carrier individuals’ phenotypic properties (Supplementary Figure 1). Here, we show the result of a query for the *de novo* LGD variants that fall in the FMRP target gene set and occur in children diagnosed with autism. The panel shows four of the 175 variants that meet all these criteria (see Figure 1d for a description of the columns). The FMRP targets are the genes that encode mRNAs targeted by the Fragile X mental retardation protein [32]. We and others have reported that FMRP target genes are enriched for damaging *de novo* variants in autism and other neurodevelopmental disorders [22]. (b) The enrichment tool (Supplementary Figure 2) allows the user to test for enrichment of *de novo* variants within a gene set. The enrichment tool results for *de novo* mutations are shown in children with autism and unaffected children in the FMRP target gene set. The top table shows that among 6,816 individuals (1,117 female and 5,699 male) with autism, 855 *de novo* LGD mutations are found (N). Of these, 175 fall in FMRP target genes (O), while we expect to see only 83 (E). This corresponds to an enrichment with a significant p-value (pV), indicated with a red background. Missense mutations are similarly enriched in FMRP target genes. However, synonymous mutations are not enriched (white background). The bottom table shows the same analysis for unaffected individuals, which finds no enrichment for LGD, missense, or synonymous mutations. (c) The phenotype browser allows users to explore the available phenotypic data associated with a dataset and download the subsets of interest. The phenotypic data within GPF is organized by *instruments,* or diagnostic tools applied to the subset of the dataset’s individuals. Each instrument includes several measures. The value of a measure can be a number, for example, for measures of IQ or height, a Yes or No answer to a question, or a free-text form. Using GPF, the user can find the list of instruments in the dataset, see all the measures for each instrument, and search for measures based on their names and descriptions. GPF provides summary-level information for each measure, including the type of its values and histograms for the values, and a mechanism to download the values for each individual for one or more measures of interest. The panel shows a part of the search results for phenotypic measures related to ‘communication’ in the SSC dataset. The results are organized as a table with one row per measure. Four measures from four different instruments are shown: the communications_standard measure from the vineland_ii instrument, the ados_communiction_social measure from the ssc_core_descriptive instruments, the communication measure from the srs_parent, and, finally, the communication measure from the srs_teachers instruments. For each measure, the table includes histograms of its values, separately by groups of individuals by gender, role, and affected status. The zoomed-in view (a feature provided by GPF) of the histograms plot for the communications_standard measure from vineland_ii shows the number of individuals for which the measure is available for the affected probands and the unaffected siblings separately by males and females and the histograms of the four groups (male and female probands and male and female siblings). The histograms show that the affected probands have diminished vineland_ii communication scores. The result table also shows the scatter plots of each of the four measures against the individuals’ ages at assessment and the individuals’ IQs. The regression lines of the measures against age and IQ are also shown. (d) The phenotype tool allows the user to test if a given phenotypic measure (e.g., non-verbal IQ) is different between the children that carry a specified type of genetic variant (e.g., *de novo* LGDs) and the children that do not have such variants (Supplementary Figure 3). The panel shows the effect of four *de novo* variant types (LGD, missense, CNV+ or large duplications, and CNV- or large deletion) in genes with low ExAC pLI rank (less than 1000) on non-verbal IQ (nvIQ). For each *de novo* variant type, the mean and the 95% confidence interval are plotted for four groups of individuals, males and females, with and without a *de novo* variant in the selected genes. There is a significant decrease in non-verbal IQ for affected children with *de novo* LGDs or CNV-within the genes with low ExAC pLI rank. There is no effect of the CNV+ mutations and only a marginal effect of missense variants.

#### Enrichment Tool

The Enrichment Tool allows the user to test if a given set of genes is affected by more or fewer *de novo* mutations in the children in the dataset than expected. We and others have used such an approach to demonstrate a functional convergence of *de novo* mutations in autism. For example, prior studies show that damaging *de novo* mutations in autistic children target synaptic genes and genes encoding chromatin modifiers [21, 23, 36]. The approach has also been used to demonstrate that the *de novo* mutations in autism target similar genes as the *de novo* mutations in schizophrenia, intellectual disability, and epilepsy [21]. Users can use the Enrichment Tool to explore their research-driven hypotheses using the extensive genetic data managed by GPF-SFARI.

To use the Enrichment Tools, a user must choose a set of genes either by selecting one of the gene sets that have already been loaded in GPF or by providing their own gene set. In addition, the user must select the dataset that contains the *de novo* variants to be used in the enrichment analysis. Finally, the user must select among the background models that GPF uses to compute the expected number of *de novo* mutations within the given dataset. GPF supports several background models, including mutability models based on sequence context [37] and models based on gene length or the number of variants of no consequence (i.e., *de novo* or rare synonymous) (Supplementary Figure 2).

Figure 3b shows an example output for the Enrichment Tool for the gene sets of the FMRP targets identified by Darnell et al [32] and *de novo* variants from the SSC dataset. The *de novo* LGDs and missense mutations from autistic children are significantly enriched in FMRP genes. In contrast, synonymous mutations are not enriched, and none of these variant types are enriched in unaffected children. These observations suggest an etiological role of the FMRP gene set in autism.

#### Phenotype browser

The Phenotype browser allows the exploration, search, and download of the available phenotypic data for the various datasets. The users can easily see the available instruments for a dataset and the phenotypic measures in each instrument. They can also search for measures across all instruments using their names and descriptions. The Phenotype browser shows the high-level statistics for each measure, including the number of individuals with values for the measure (or the number of individuals for which the instrument has been applied) and histograms of the measures’ values, separately for different roles (i.e., proband, mother, sibling), affected statuses, and genders. It can also provide correlations of all measures with a set of core phenotypic properties. For example, the Phenotype browser for the SSC dataset shows the correlations of all measures with the individual’s age and IQ. Figure 3c shows the high-level statistics for four measures related to ‘communication’ in the SSC dataset.

#### Phenotype/Genotype tool

The “Phenotype/Genotype Tool” is a powerful resource that enables users to explore associations between genotypes and phenotypes, harnessing the wealth of integrated data from both domains (Figure 3d). Its operation is straightforward: users select a specific phenotypic attribute and choose a type of genetic variant based on frequency and the effect on target genes. Subsequently, the tool assesses whether individuals carrying such a variant exhibit statistically significant differences in the chosen phenotypic attribute when compared to individuals lacking that variant (Supplementary Figure 3). This tool has played a pivotal role in our research and published findings. For instance, we showed that children with autism who harbor detrimental *de novo* mutations tend to display lower IQ levels [21, 25] and impaired motor skills in comparison to their counterparts without such variants [38, 39].

### GPF “at home”

GPF is an open-source project with comprehensive documentation (https://iossifovlab.com/gpfuserdocs/) that others can easily install and use to manage their genotypic and phenotypic data. It can be used on small datasets (as small as data for one family) but scales to datasets with genotypes and phenotypes from hundreds of thousands or even millions of individuals. Users can seamlessly browse, analyze, and securely share their data with colleagues and the broader scientific community (Figure 4).

**Figure 4.**
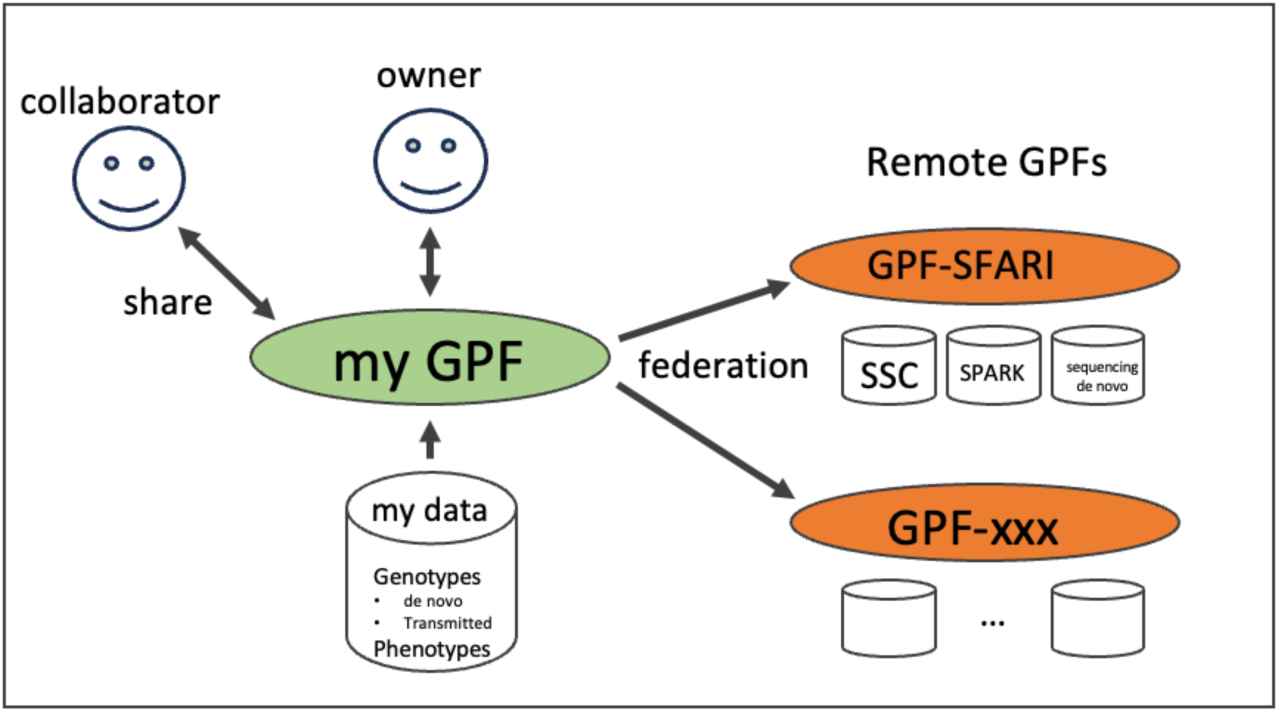
GPF at home. Users can instantiate their own GPF (my GPF) to manage and analyze their data. Through GPF, they may also share their data with other researchers. Finally, users may connect to other GPF instances (e.g., GPF-SFARI) using the Federation feature and perform joint queries and analysis of their data and the data in the remote instances.

The “Federated access” feature extends GPF’s capabilities by allowing one GPF instance to connect with one or more additional GPF instances. Through federated access, users can jointly analyze the data in multiple GPF instances. A common scenario involves a user configuring a GPF instance to work with their private data while accessing a public GPF instance (like the one at SFARI) hosting relevant datasets and resources. Once the necessary authorization is obtained for remote GPF instance data, establishing federated access within the local GPF instance is straightforward.

## Discussion

Here, we described our platform, GPF, which offers unparalleled versatility for exploring and analyzing genetic and phenotypic variation in families. GPF accommodates the genetic variant and family pedigree types typically used in genetic analysis and enables users to browse and select variants using an extensive array of variant properties and phenotypic measures of the carrier individuals. The GPF power tools allow the user to conduct enrichment and phenotypic association analysis. As a flexible and scalable platform capable of handling genotypic and phenotypic data from millions of individuals, GPF can be applied to a large class of genetic study types.

We used GPF on the extensive autism genetic and phenotypic data collected by SFARI, corresponding to ∼9,000 and ∼300,000 individuals in the SSC and SPARK collections. The resulting variant collections, as well as the associated phenotypic measures, are accessible through our GPF instance, GPF-SFARI. While these two data sets are accessible only after registration, a third one, Sequencing *de novo*, which includes *de novo* variants from studies of six developmental disorders, is publicly available at GPF-SFARI. GPF-SFARI also provides a freely accessible collection, Gene Profile View, of statistics for each gene regarding the number of genetic variants of different types in the three datasets without any connection to sensitive individual-level data. The powerful interactive query interface and the analysis tools of GPF enable researchers to use the rich data set of genotypic and phenotypic variation in GPF-SFARI easily and effectively.

## Methods

### Architecture and implementation

The structure of GPF comprises three stacked layers (Supplementary Figure 4). The system’s core is the Data Access Environment (DAE) layer. The DAE is implemented in Python 3 and provides access to all underlying genotypic and phenotypic data, all relevant genomic information (i.e., the reference genome, gene, and variant properties), the set of authorized users and their access privileges (represented as user groups), and analysis tools.

The phenotypic and genotypic data are stored in Phenotype and Genotype storages, respectively. The Phenotype Storage is implemented as a relational data schema that can be managed by various relational database systems, like SQLite (https://www.sqlite.org) or MySQL (https://www.mysql.com). We based our Genotype Storage on the modern columnar database organization to enable interactive access even for a genotyping data set over millions of individuals. GPF’s Genotype storages can be deployed on different frameworks. The DuckDB (https://duckdb.org) Genotype Storage is useful for small to intermediately sized datasets and requires no management. The Impala (https://impala.apache.org) Genotype Storage is a distributed server environment that can handle any data size and be set up on an existing computational infrastructure. The BigQuery (https://cloud.google.com/bigquery) Genotype Storage operates on Google’s cloud infrastructure, requires minimal management, and can handle any data size. The BigQuery’s drawback is that it may prove expensive for systems with high loads.

The second layer in the GPF is the WDAE REST API. It uses the DAE layer and is implemented based on Python’s Django framework (https://www.djangoproject.com). WDAE allows remote scripts to access the data and analysis tools the DAE provides. The last layer is the GPFs’s user web interface. It is implemented on top of the WDAE layer with the help of the AngularJS JavaScript framework (https://angularjs.org).

The three layers enable users to interact with GPF in different ways. The GPF web interface enables all users to access the data interactively. Everyone can also download the relevant data for further analysis, usually in Excel table form. Users with a computational background can also create scripts to automate the interaction with the system. For example, if a researcher uses Python and has access to the computer where GPF is deployed, they can create Python scripts to access the data through the DAE API. Alternatively, a researcher who prefers other programming languages or doesn’t have access to the GPF’s host can still create scripts accessing the GPF’s data and tools through the WDAE interface.

### Annotation

Genotypic data analyses rely on examining the genotypes in the context of the vast amounts of genomic data generated in recent decades. To facilitate joint analysis of multiple datasets, GPF’s import process annotates all the variants uniformly across all imported studies with a user-specified list of genomic properties. The source for these properties is one or more Genomic Resource Repositories (GRR). We have compiled a sizeable GRR and made it accessible at https://storage.googleapis.com/iossifovlab-grr/index.html. Our repository comprises publicly available resources, including population frequencies from Gnomad [1], conservation scores like CADD [13], phyloP [10], and phastCons [11], and missense scores like MPC [12]. Each resource within the repository is assigned a resource identifier. GPF users only need to specify the list of resource identifiers that they wish to annotate the GPF instance variants with. GPF’s GRR infrastructure is flexible and allows users to configure one or more private GRRs, including other public or private resources, and annotate their variants with the additional resources.

### Import and Import Cluster

DAE supports importing data in standard file formats where appropriate, for example, pedigree (.ped) files to describe families. When the standard formats were unavailable or could not meet our requirements, we defined our custom formats, like the file format for phenotypic data.

GPF can import genotypes from two main file formats: the standard VCF format [40] and a simple list of variants typically used for importing *de novo* variants. The GPF’s import functions are flexible and can directly handle a variety of input file configurations. For example, a researcher can equally easily import a single VCF file with all his data, multiple VCF files, each containing genotypes for a human chromosome, or multiple VCF files, each representing genotypes for a group of families.

The phenotypic data is imported from a set of simple files (one for each instrument) organized as a table with columns representing the instrument’s measures and lines representing the individual to which the instrument has been applied.

Importing a large dataset in a GPF instance is slow, particularly if the user has requested many annotation properties. GPF allows the user to configure an ‘Import Cluster’ that he can use to parallelize and thus substantially speed up the import process. The ‘Import Cluster’ can be configured to run on multiple cores of one host, a local computation cluster like clusters controlled by OGE (https://www.oracle.com/technetwork/oem/host-server-mgmt/twp-gridengine-overview-167117.pdf) or Slurm (https://slurm.schedmd.com/), or a cloud Kubernetes (https://kubernetes.io) cluster.

## Acknowledgments

This work was supported by the Simons Center for Quantitative Biology at Cold Spring Harbor Laboratory, SFARI Grants SF497800, SF529232, SF677963, SF666590 (to I.I.), the Centers for Common Disease Genomics grant (UM1 HG008901). The National Human Genome Research Institute and the National Heart, Lung, and Blood Institute fund the Centers for Common Disease Genomics.

## Supplementary Figures

**Supplementary Figure 1.**
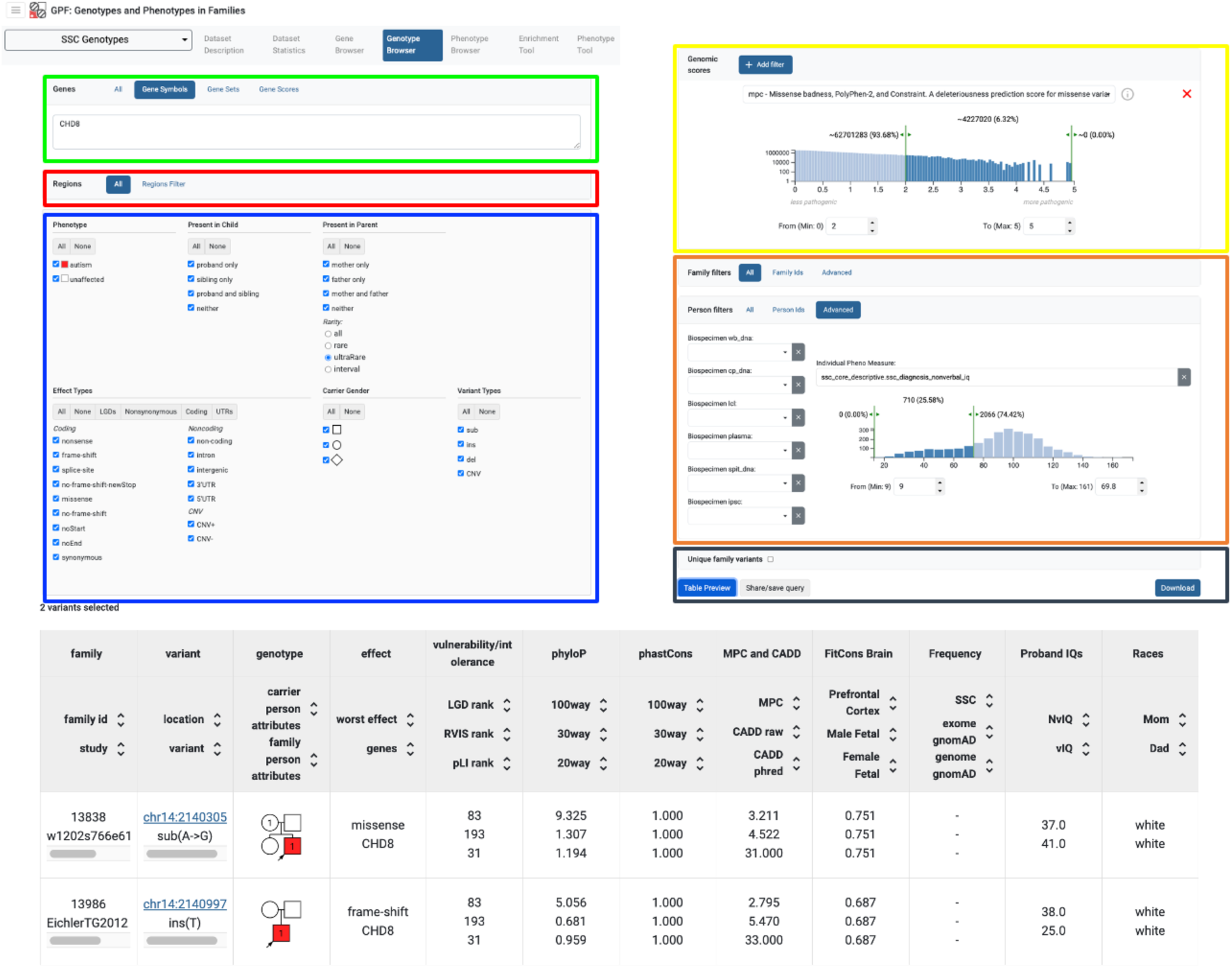
Genotype browser. The Genotype Browser tab allows users to access the genetic variants associated with a particular dataset. The figure shows one example query and its results displayed in the Genotype Browser tab for the “SSC Genotypes” dataset. The user interacts with the Genotype Browser by first specifying the filters on the properties of the genetic variant of interest and then pressing one of the result buttons (black rectangle), “Table Preview” or “Download,” to receive the results into the desired form. The filters of the various types of properties are arranged in separate panels above the result buttons. In the Genes panel (green rectangle), the users can specify the genes or gene priorities (i.e., “Gene Sets” or “Gene Weights”) that are affected by the genetic variants. In the Regions panel (red rectangle), the user can specify the genomic regions of interest. In the next panel (blue rectangle), the user can specify various constraints on the properties of the genetic variant, including the type of variant (i.e. SNVs or CNVs), the effect of the variant on the targeted genes (i.e. missense or synonymous) and the transmission pattern (i.e. present in mother, in father, or de novo, present in affected in unaffected children). In the Genomic Scores panel (yellow rectangle), the user can specify constraints on various genomic scores associated with variants (i.e., CADD, polyPhen, or MPC). Finally, in the Families and Person filters panels (orange rectangle), the user can specify the families or individuals of interest or constraints on the phenotypic properties (i.e., non-verbal IQ) associated with individuals carrying the genetic variants. In this example, the user has specified that they are interested only in variants affecting the *CHD8* gene that are ultra-rare (defined as seen only once in the dataset), that have an MPC score larger than 2, and are present in individuals with non-verbal IQ less than 70. The user has pressed the “Tabular Preview” button, which has resulted in the 2 genetic variants displayed in a table form below the results buttons. Each row of the table describes one variant segregating in one family. The first four columns specify the core properties of the variant, the family where the variant segregates, the definition of the sequence variant (its location and the nucleotide sequence change), the affected gene, and the effect the variant has on the genes. The next columns include properties of the targeted genes (vulnerability scores), properties of the variant itself (the phyloP, phastCons, MPC, CADD, and FitCons brain genomic scores and variant frequencies from different cohorts), and phenotypic properties of the carrier individuals (verbal and non-verbal IQs and father’s and mother’s races).

**Supplementary Figure 2.**
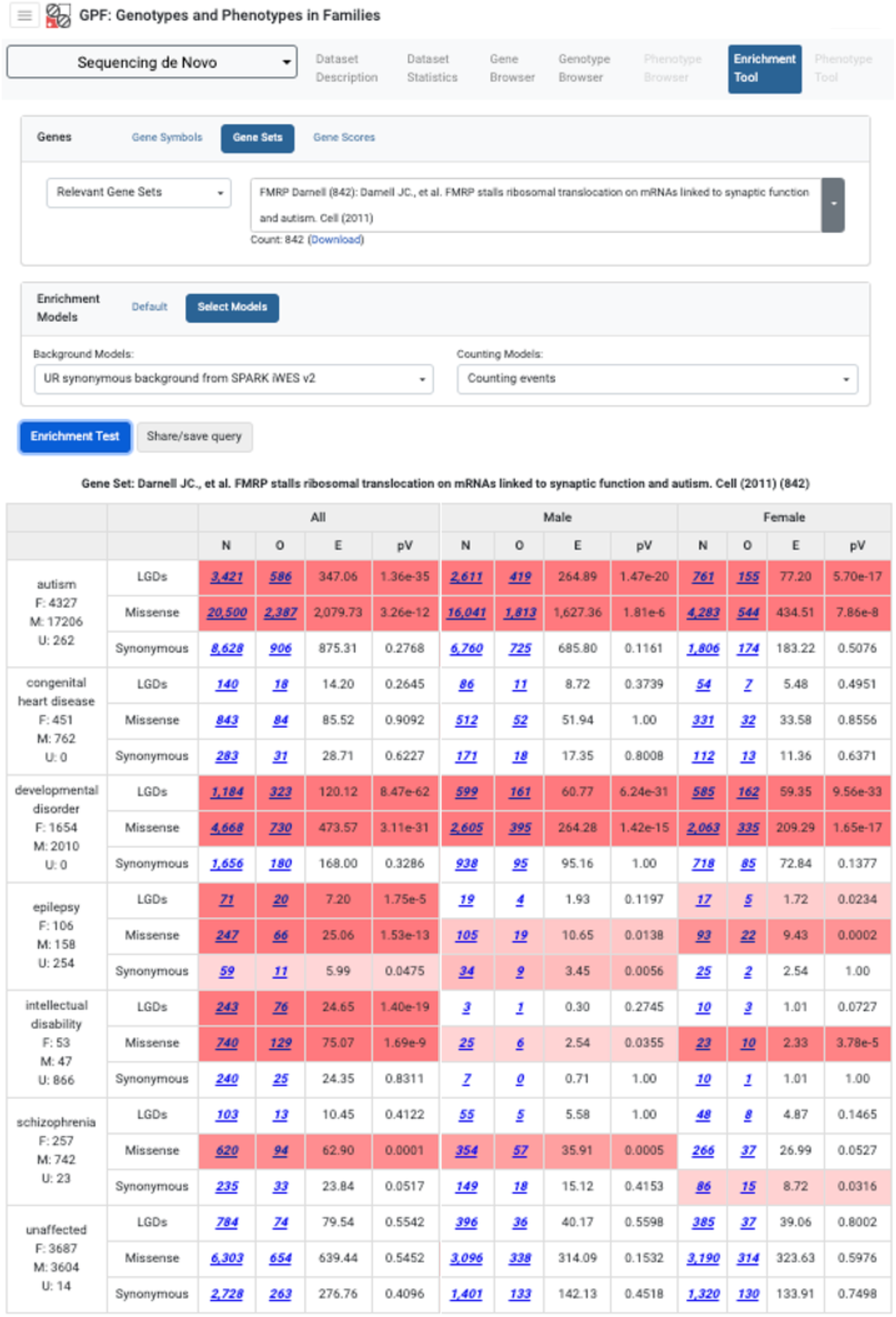
Enrichment. The figure shows an example of using the Enrichment Tool. The Enrichment Tool allows the user to test if a given set of genes is affected by *de novo* mutations more or less than expected in the children in the selected dataset. The user selects the genes to be tested using the Genes panel (green rectangle) and computes expected numbers of *de novo* mutations of three types, LGD, missense, and synonymous, in groups of children from the datasets defined by the primary diagnosis. The Enrichment Tool displays the total number of variants of the particular type for each diagnosis (N columns), the observed number of these variants that affect a gene from the selected genes (O columns), and the expected number of variants in the genes (E). The expected number of variants is computed using a background model that the user can control through the Enrichment Models panel. Finally, the tool computes and displays a p-value (pV columns) for the observed number of variants in the genes based on a Poisson distribution with a parameter equal to the expected number of variants in the selected genes. The example above shows the enrichment results in the Sequencing de novo Dataset that includes children with six primary diagnoses (autism, congenital heart disease, epilepsy, intellectual disability, schizophrenia, developmental disorder, and unaffected control children) for the set of FMRP target genes [32]. A significant enrichment (red background) is seen for non-synonymous variants in autism, epilepsy, intellectual disability, and schizophrenia but not in congenital heart disease or of the unaffected controls. No synonymous variants in the FMRP target genes are enriched in any phenotype.

**Supplementary Figure 3.**
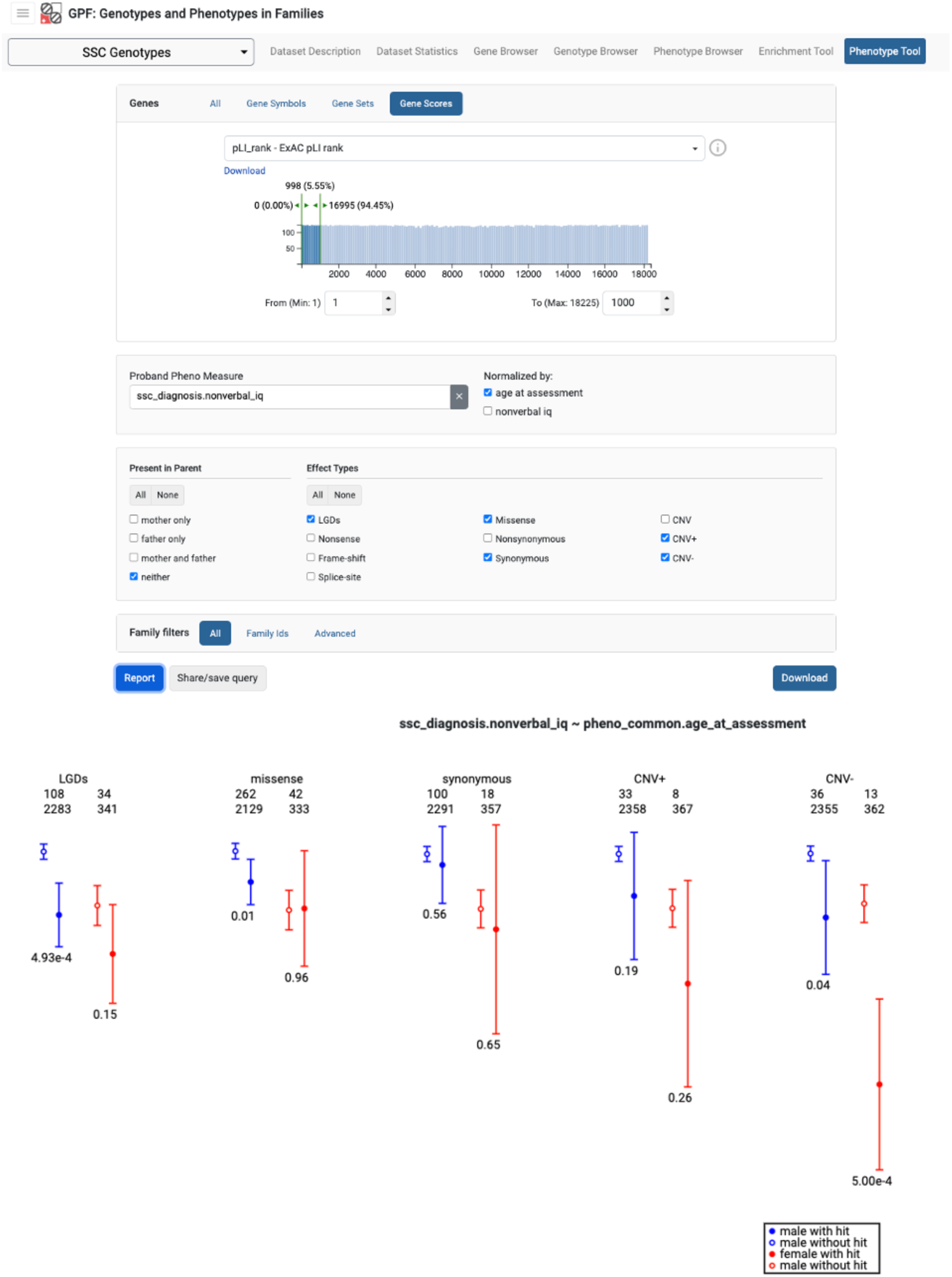
Phenotype/Genotype Tool. The Phenotype/Genotype Tool tests if a given phenotypic measure (i.e., non-verbal IQ) is different between the children that carry the specified type of genetic variant (i.e., *de novo* LGDs) and the children that do not have such variants. The example shows the effect on non-verbal IQ (the nonverbal_iq measure from the ssc_diagnosis instrument, ssc_diagnosis.nonverbal_iq) of the variants in genes with pLI rank of less than or equal to 1000, or the 1000 genes most intolerant to loss-of-function mutations. For example, we see a significant decrease in non-verbal IQ for affected males with de novo LGDs and females with a large de novo deletion (CNV-) affecting one of the selected genes (both with p-value ∼ 5*10^−4^). In addition, large *de novo* duplications (CNV+) exhibit no statistically significant impact in both genders, while the influence of *de novo* missense variants is only marginal in males (p-value = 0.01).

**Supplementary Figure 4.**
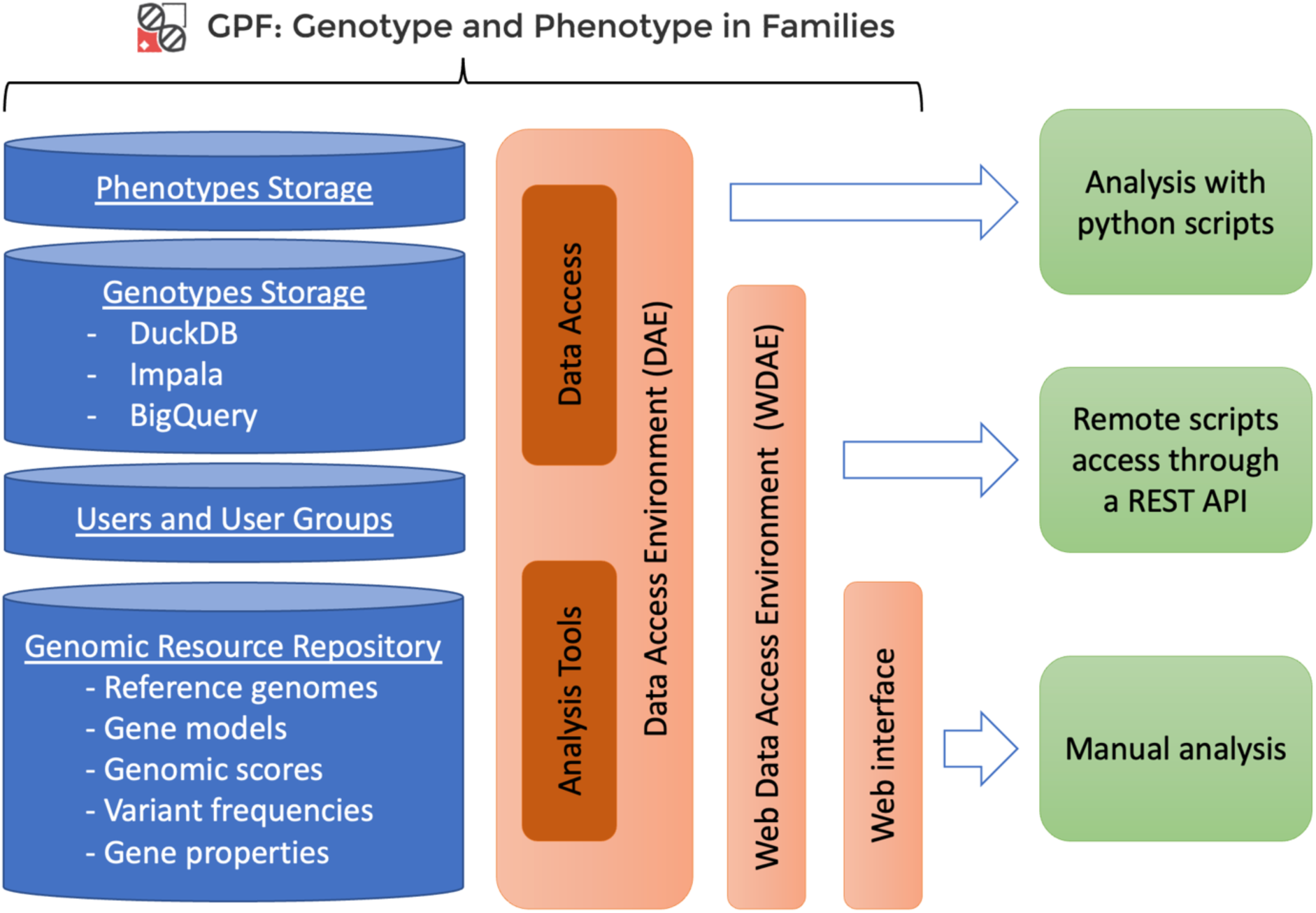
GPF Architecture. GPF’s architecture consists of three integrated layers. At its core is the Data Access Environment (DAE) layer, serving as the system’s foundation, with access to Phenotype, genotype, user, and genomic resource repository information. The second layer introduces the WDAE REST API, enabling remote scripts to interface with the data and analysis tools the DAE offers. The outermost layer is GPF’s user web interface, allowing users to execute intricate queries and utilize the power tools outlined in the manuscript.

